# Development and validation of Arc nanobodies: new tools for probing Arc dynamics and function

**DOI:** 10.1101/2022.01.20.477070

**Authors:** Yuta Ishizuka, Tadiwos F. Mergiya, Rodolfo Baldinotti, Ju Xu, Erik I. Hallin, Sigurbjörn Markússon, Petri Kursula, Clive R. Bramham

**Affiliations:** Department of Biomedicine, University of Bergen, Bergen, Norway; Mohn Center for Research on the Brain, University of Bergen, Norway; Faculty of Biochemistry and Molecular Medicine, University of Oulu, Oulu, Finland; Biocenter Oulu, University of Oulu, Finland

**Keywords:** Arc, N-lobe, C-lobe, nanobody, intrabody, chromobody, co-immunoprecipitation

## Abstract

Activity-regulated cytoskeleton-associated (Arc) protein plays key roles in long-term synaptic plasticity, memory, and cognitive flexibility. However, an integral understanding of Arc mechanisms is lacking. Arc is proposed to function as an interaction hub in neuronal dendrites and the nucleus, yet Arc can also form retrovirus-like capsids with proposed roles in intercellular communication. Here, we sought to develop anti-Arc nanobodies (ArcNbs) as new tools for probing Arc dynamics and function. Six ArcNbs representing different clonal lines were selected from immunized alpaca. Immunoblotting with recombinant ArcNbs fused to a small ALFA-epitope tag demonstrated binding to recombinant Arc as well as endogenous Arc from rat cortical tissue. ALFA-ArcNb also provided efficient immunoprecipitation of stimulus-induced Arc after carbachol-treatment of SH-SY5Y neuroblastoma cells and induction of long-term potentiation in the rat dentate gyrus *in vivo*. Epitope mapping showed that all Nbs recognize the Arc C-terminal region containing the retroviral Gag capsid homology domain, comprised of tandem N-and C-lobes. ArcNbs E5 and H11 selectively bound the N-lobe, which harbors a peptide ligand binding pocket specific to mammals. Four additional ArcNbs bound the region containing the C-lobe and terminal tail. For use as genetically encoded fluorescent intrabodies, we show that ArcNbs fused to mScarlet-I are uniformly expressed, without aggregation, in the cytoplasm and nucleus of HEK293FT cells. Finally, mScarlet-I-ArcNb H11 expressed as intrabody selectively bound the N-lobe and enabled co-immunoprecipitation of full-length intracellular Arc. ArcNbs are versatile tools for live-cell labeling and purification of Arc and analysis of capsid domain specific functions.

## 1. Introduction

The immediate early gene encoding activity-regulated cytoskeleton-associated protein (Arc), also known as activity-regulated gene 3.1 (Arg3.1), is expressed mainly in excitatory neurons following neural activity to mediate functionally diverse actions [1]. In activity-dependent synaptic plasticity, such as long-term potentiation (LTP) and long-term depression, Arc protein is rapidly synthesized and degraded, indicating a transient and dynamic mode of action [2-5]. Stimulus-evoked expression of Arc is also required for memory, postnatal developmental plasticity of the visual cortex, and cognitive flexibility [3, 6-9]. These diverse functions are proposed to be mediated by distinct Arc-binding partner complexes in the postsynaptic compartment and the neuronal nucleus [10-14]. Arc protein is also capable for reversible self-association and formation of oligomers [15, 16]. Recently, Arc protein was shown to self-assemble into spheroid particles resembling retroviral Gag capsids [17-19]. Arc capsids are incorporated in extracellular vesicles and are capable of transferring RNA cargo from donor cells to recipient cells [17, 18]. However, an integral understanding that accounts for Arc hub signaling and capsid function is lacking.

Arc is a flexible protein comprised of two oppositely charged domains, N-and C-terminal domains (NTD and CTD), flanking a central disordered hinge region [15]. Structural studies and predictions indicate an antiparallel coiled-coil structure of the NTD [20]. The NTD mediates binding to phospholipid membrane [20, 21], and the NTD second coil harbors an oligomerization motif ^113^MHVWREV^119^ that is critical for Arc self-association including formation of higher-order oligomers and capsids, but is not required for dimer formation [19]. The CTD is a tandem domain comprised of two lobes, the N-and C-lobes (NL and CL), homologous to the capsid domain of retroviral Gag polyprotein [22, 23]. Arc CTD is important for formation of stable and double-shelled Arc capsid, which is dependent on NTD-mediated oligomerization [18, 19, 24]. In addition, the mammalian Arc NL has a unique hydrophobic peptide binding pocket that interacts with several postsynaptic proteins, including stargazin, guanylate kinase-associated protein (GKAP), Wiskott-Aldrich syndrome protein family member 1 (WAVE1), and GluN2A [14, 20, 22, 25]. Taken together, this suggests that interactions between domains regulates oligomerization and protein-protein interactions to determine Arc activity-state. However, tools that would allow labeling or manipulation of endogenous Arc in situ are lacking.

Single-domain antibodies (sdAb), also known as variable domain of heavy chain of heavy chain (VHH) antibody and Nanobody® (Nb), are derived from the heavy chain antibody of *Camelidae* [26]. Nbs have unique properties relative to conventional antibodies that make them attractive for use as research tools and development of therapeutics [27]. Nbs are small (15 kDa), have high chemical stability, can be genetically-encoded for expression in mammalian cells as intrabodies, and they can bind hidden epitopes in small cavities that are not recognized by antibodies [28-32].

Here, we sought to develop and validate experimental applications for six anti-Arc nanobodies (ArcNbs: B5, B12, C11, D4, E5, and H11). ArcNbs fused to a small ALFA epitope were successfully used for immunoblotting and immunoprecipitation of endogenous Arc. ArcNb E5 and H11 specifically recognized the capsid domain NL, while the other Nbs bound the segment containing the CL and C-terminal tail. Genetically-encoded ArcNb fused to fluorescent mScarlet-I expressed efficiently in mammalian cells as intrabodies and allowed immunoprecipitation of intracellular Arc. The ArcNbs provide versatile tools for labeling, tracking, and purifying native intracellular Arc, with further potential use as capsid domain-specific inhibitors.

## 2. Materials and Methods

### 2.1. Generation and modifications of ArcNbs

Procedures for generation and modification of ArcNbs were conducted by NanoTag Biotechnologies GmbH (Göttingen, Germany) (**Fig. 1**). For detailed information on the purification, structural characterization, and binding of the untagged ArcNbs, see Markússon et al. (2021). Briefly, two alpacas were immunized six times starting with purified recombinant human wild-type (WT) Arc protein (immunization 1 and 2), then a mixture of Arc WT and purified mutant Arc protein (Arc^s113-119A^: amino acid residues from 113 to 119 were substituted to alanine) with increasing proportion of Arc^s113-119A^ (immunization 3-5), and finally with injection of Arc^s113-119A^ alone. A strong general immune-response was confirmed using a serum-based ELISA in one of the animals. Total RNA was extracted from peripheral blood mononuclear cell (PBMC) preparations obtained from whole blood. cDNA encoding Nbs were reverse transcribed using a nested PCR that specifically amplifies the coding regions for IgG2 and IgG3 VHH fragments. The complete PCR product was cloned into a phagemid vector suitable for phage display followed by bio-panning with purified mutant Arc^s113-119A^ protein as antigen. Then, Nb sequences extracted from phages binding to antigen were amplified by PCR and cloned in the screening vector. Ninety-six Nb clones were separately expressed in *E. coli*. and validated by ELISA using Arc^s113-119A^ as target protein. Six clones (B5, B12, C11, D4, E5, and H11) were chosen based on their sequences and ELISA-based binding evaluations.

**Fig. 1.**
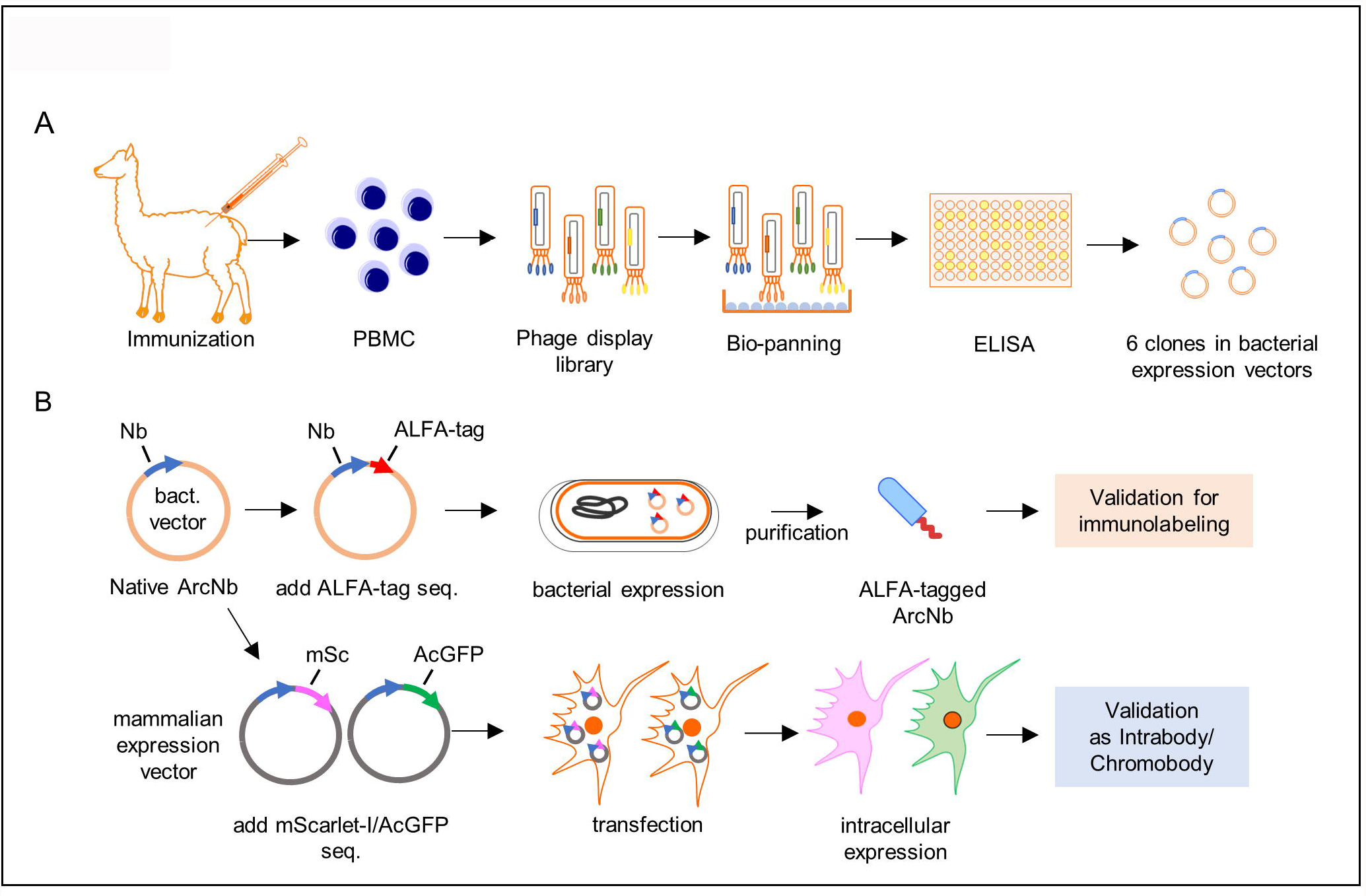
Schematic of ArcNb generation and modification. ***A***, Schematic of ArcNb generation pipeline. ***B***, Modifications of ArcNb for biochemical and immunolabeling applications. Detailed information is described in **Materials and Methods** section.

To generate recombinant Nbs suitable for biochemical analyses, such as immunoblotting and immunoprecipitation, a small ALFA-epitope tag (SRLEEELRRRLTE) [33], which forms a stable alpha-helix on target protein, was added to the Nb C-terminus. For expressing ArcNbs as intrabody/chromobody, the Nb coding regions were subcloned from the bacterial expression vector into a mammalian expression vector harboring mScarlet-I (mSc) driven by the CMV promoter. We also subcloned the coding region of all ArcNbs into the AcGFP-N1 vector. AcGFP1-N1 was a gift from Michael Davidson (Addgene plasmid # 54705; http://n2t.net/addgene:54705; RRID:Addgene 54705).

### 2.2. Purification of ALFA-tagged ArcNbs

ALFA-tagged ArcNbs (ALFA-ArcNbs) were expressed in BL21 (DE3) cells and purified according to a general methods for protein purification [34] with minor modifications. Briefly, cells were grown at 37°C in LB medium overnight. The cultures were transferred to ZY5052 medium for auto-induction of protein expression at 37°C for 4 hours, then at 30°C for 24 hours. The cells were lysed in lysis/washing buffer containing 40 mM Hepes (pH 7.5), 100 mM NaCl, and 20 mM Imidazole, by one freeze-thaw cycle followed by sonication. The lysates were centrifuged at 16,000 x *g* for 30 min at 4°C and loaded onto a Ni-NTA resin. After washing resin with lysis/washing buffer, ALFA-ArcNbs were eluted by elution buffer containing 40 mM Hepes (pH 7.5), 100 mM NaCl, and 300 mM Imidazole. His-tagged TEV protease was added to the elute, and the samples were dialyzed against dialysis buffer containing 40 mM Hepes (pH 7.5), 100 mM NaCl and 1 mM dithiothreitol (DTT) overnight at 4°C. The dialyzed samples were passed through a Ni-NTA resin again to remove the TEV protease and the cleaved His10-tag. The Ni-NTA flow-through was loaded on a chromatography column (HiLoad® 16/60 Superdex® 75 PG, Cytiva, Marlborough, MA), equilibrated with degassed buffer (20 mM Tris-HCl pH 7.4, 150 mM NaCl). Size exclusion chromatography was run at 1 ml/min at 4°C on a ÄKTA pure chromatography system (Cytiva). All ALFA-ArcNbs gave one major peak in the chromatogram. The protein size and purity were analyzed by SDS-PAGE, giving one strong Coomassie-stained band of the expected size (**Fig. S1**).

### 2.3. Animals, electrophysiology, and tissue collection

For naive cortical tissue collection, adult male Sprague-Dawley rats were anesthetized using urethane (1.5 g/kg intraperitoneal) and decapitated before dissecting out cortical tissues. The tissues were kept in −80°C until experiments. LTP in the dentate gyrus (DG) was induced according to previously described methods [35]. Briefly, a bipolar stimulating electrode (7.9 mm posterior to the bregma, 4.2 mm lateral to the midline) and recording electrode (3.9 mm posterior to bregma, 2.3 mm lateral to midline) were inserted to stimulate medial perforant path fibers and record of evoked field potentials in the dentate hilus, respectively. Test pulses were given every 30 s throughout the experiment except during the period of high-frequency stimulation (HFS). 20 min baseline recordings were acquired before three sessions of HFS with five min interval were given. Each session consisted of four, 400 Hz stimulus trains (8 pulses/train) and the interval between trains was 10 s. Changes in the field excitatory post-synaptic potential slope were expressed in percent of baseline to quantify induction and maintenance of LTP. Two hours after HFS, rats were decapitated, and the dentate gyri were dissected out and stored at -80°C for further use.

### 2.4. Cell culture: transfection and pharmacological stimulation

Human embryonic kidney 293FT (HEK293FT) and SH-SY5Y neuroblastoma cells were obtained from Thermo Fisher Scientific (Waltham, MA) and ATCC (Manassas, VA), respectively. Cells were maintained in Dulbecco’s modified Eagle medium (DMEM/high-glucose, Sigma-Aldrich, St. Louis, MO) supplied with 10% FBS (Sigma-Aldrich) and 100 U/ml Penicilin-Streptomycin (Thermo Fisher Scientific) at 37 °C in a humidified incubator with 5% CO_2_. HEK293FT cells were transfected using Lipofectamine™ 2000 Transfection Reagent (Thermo Fisher Scientific) according to the manufacturer’s instructions. For induction of endogenous Arc expression, SH-SY5Y cells were stimulated with 100 μM carbachol (CCh) for 1 h at +37 °C in a humidified incubator with 5% CO_2_ [36].

### 2.5. Antibodies

Primary antibodies are as follows: mouse anti-Arc (C-7) (Santa Cruz Biotechnology, Dallas, TX, Cat# sc-17839, RRID:AB_626696), rabbit anti-Arc (Synaptic Systems, Göettingen, Germany, Cat# 156 003, RRID:AB_887694), rabbit anti-GFP (Santa Cruz Biotechnology Cat# sc-8334, RRID:AB_641123), mouse anti-mCherry (Takara Bio, Siga, Japan, Cat# 632543, RRID:AB_2307319), anti-ALFA HRP-coupled sdAb (NanoTag Biotechnologies, Cat#. N1501-HRP). Secondary antibodies are as follows: Goat Anti-Mouse IgG, H & L Chain Antibody, Peroxidase Conjugated (Merck, Darmstadt, Germany, Cat# 401253, RRID:AB_437779) and Goat Anti-Rabbit IgG, H & L Chain Specific Peroxidase Conjugate antibody (Merck Cat# 401315, RRID:AB_2617117). Secondary antibodies for detecting immunoprecipitated proteins are as follows: Peroxidase AffiniPure Goat Anti-Mouse IgG, light chain specific (Jackson ImmunoResearch Labs, West Grove, PA, Cat# 115-035-174, RRID:AB_2338512) and Peroxidase IgG Fraction Monoclonal Mouse Anti-Rabbit IgG, light chain specific (Jackson ImmunoResearch Labs, Cat# 211-032-171, RRID:AB_2339149).

### 2.6. Electrophoresis and immunoblotting

Native-polyacrylamide gel electrophoresis (PAGE) was performed according to previously described methods [37]. Briefly, different amounts of purified mutant Arc^s113-119A^ protein were dissolved in native sample buffer, then subjected to nativePAGE using 4−15% Tris-Glycine acrylamide gel. SDS-PAGE was performed as described previously [38, 39]. Briefly, cultured cells and brain tissues were lysed and sonicated in RIPA buffer containing cOmplete™, EDTA-free Protease Inhibitor Cocktail Tablet (Sigma-Aldrich) and PhosSTOP™, Phosphatase Inhibitor Cocktail Tablet (Sigma-Aldrich). After centrifugation, the supernatants were collected, and protein concentration was determined using the Micro BCA Protein Assay Kit (Thermo Fisher Scientific). Samples were mixed with 4X Laemmli sample buffer (Bio-Rad Laboratories, Inc., Hercules, CA) containing 50 mM DTT at final concentration, then boiled and subjected to SDS-PAGE. Separated proteins in the gel then were transferred to polyvinylidene difluoride membrane (Thermo Fisher Scientific). The membranes were incubated with the appropriate primary and secondary antibodies after incubation with 10% bovine serum albumin (BSA) in Tris-buffered saline with 0.05% Tween 20 (TBS-T) at room temperature for 1 h. For detecting Arc protein immunolabeled by ArcNb, anti-ALFA HRP-coupled sdAb was used instead of conventional secondary antibody. Peroxidase activity was detected using chemiluminescence regent (Clarity Western ECL substrate, Bio-Rad Laboratories, Inc.) and visualized by an image analyzer using Image Lab™ Software (Gel Doc™ XR+, Bio-Rad Laboratories, Inc.). Figure presentations were performed using ImageJ/FIJI (RRID: RRID:SCR_002285).

### 2.7. Immunoprecipitation of endogenous Arc protein with ALFA-ArcNbs

Endogenous Arc immunoprecipitation was performed using ALFA-ArcNbs and ALFA Selector^ST^ as carrier [33]. CCh-treated SH-SY5Y cells and DG tissue from HFS-treated rats were lysed in modified RIPA (M-RIPA) buffer (50 mM Tris-HCl pH 7.4, 150 mM NaCl, 1% NP-40, 0.25% sodium deoxycholate, 1 mM EDTA) supplemented with cOmplete™ (Sigma-Aldrich) and PhosSTOP™ (Sigma-Aldrich). First, 20 µl of ALFA Selector^ST^ resin washed twice in 1 ml M-RIPA buffer and centrifuged at 1000 *g* for 1 min at 4 °C. Then, the resin was incubated with 2 µg of ALFA-ArcNb H11 for 1 hour at 4°C with head-over-tail rotation. Following washing, the ALFA-ArcNb H11 and selector resin complex was incubated with lysates for 30 min at room temperature. After washing resin, the samples were mixed with 2X Laemmli sample buffer containing 100 mM DTT at final concentration and denatured by boiling at 95 °C for 5 min. The supernatants were subjected to SDS-PAGE followed by immunoblotting for analyzing precipitated complexes.

### 2.8. Fluorescence cell imaging

Preparation of coverslips for cell imaging were performed according to previously described methods [39]. HEK293FT cells seeded on coverslips in 12-well plates were transfected with mSc-ArcNb expression vectors according to the manufacturer’s instructions. Transfected cells were fixed with 4% paraformaldehyde in 0.1 M phosphate buffer (pH 7.4) for 20 min at room temperature. After washing coverslips with phosphate-buffered saline (PBS, pH 7.4), the coverslips were mounted in Prolong™ Diamond Antifade Mountant with DAPI (Thermo Fisher Scientific). Fluorescence imaging was performed on AxioImager Z1 microscopy (Carl Zeiss AG, Oberkochen, Germany). Transfected cells were imaged using Plan-ApoChromat 20x/0.75 NA (Carl Zeiss AG), EC plan Neofluar 40x/1.30 Oil M27 (Carl Zeiss AG) objectives, and Axiocam 503 mono digital camera controlled by ZEN pro software (Carl Zeiss AG). The captured fluorescence images were analyzed using MetaMorph® imaging software (Molecular Devices, San Jose, CA) and ImageJ/Fiji.

### 2.9. Co-immunoprecipitation

HEK293FT cells were co-transfected with plasmids expressing mSc-ArcNb H11 and an mTurquoise2 (mTq2)-tagged Arc construct. mTq2 was N-terminally fused to Arc full-length (FL), 2) N-terminal region (NTR 1-140), 3) linker (135-216), 4) N-lobe (NL 208-277), or 5) C-lobe + C-terminal tail (CL+tail 278-396). Transfected cells were lysed in modified RIPA (M-RIPA) buffer. First, 2 μg of normal rabbit IgG (Merck, Cat# 12-370, RRID:AB_145841), anti-mCherry (mCh) antibody against mSc-ArcNb (H11), or anti-Arc rabbit polyclonal antibody were absorbed to Protein G Sepharose 4 Fast Flow (Cytiva) for 1 h at room temperature. Then, antibody/bead complexes were mixed with the cell lysates overnight at +4 °C for co-IP. The sepharose beads were washed four times with M-RIPA buffer and resuspended in 2X Laemmli sample buffer containing 100 mM DTT before denaturing at 95 °C for 5 min. The supernatants were subjected to SDS-PAGE followed by immunoblot analysis.

## 3. Results

### 3.1. ALFA-ArcNbs recognize purified recombinant Arc and endogenous Arc protein

Two alpacas were immunized with a combination of purified recombinant human WT Arc and rat mutant Arc^s113-119A^ [40]. WT Arc self-associates and oligomerizes *in vitro* [15, 16]. We have recently shown that amino acid residues 113-119 are critical for high-order oligomerization [19]. Interestingly, the substitution mutant protein Arc^s113-119A^ failed to form high-order oligomers or capsids. A mixture of WT and Arc^s113-119A^ mutant protein was used as an antigen cocktail because oligomerization of WT Arc protein might lead to epitope masking preventing proper immune response. Following bio-panning assay using Arc^s113-119A^ as a target, 96 clones were expressed and evaluated by ELISA assay using mutant Arc^s113-119A^ as an antigen. Six clones (B5, B12, C11, D4, E5, and H11) representing different clonal lines were chosen based on their sequences and ELISA-based binding to purified Arc^s113-119A^.

Firstly, we examined whether ALFA-ArcNbs have ability to detect purified recombinant Arc and endogenous Arc from rat cortical lysates. In immunoblot assays, ALFA-ArcNb bound to Arc was detected by using anti-ALFA HRP-coupled sdAb. All six ArcNbs bound to purified Arc^s113-119A^ both in native and denatured conditions (**Fig. S2**). On native-PAGE gels (**Fig. S2A**), several immunoreactive bands were detected as low-order oligomeric forms. In the denatured condition on SDS-PAGE (**Fig. S2B**), only one prominent immunoreactive band was detected around 50 kDa corresponding to Arc monomer. Also, immunoblotting performed in cortical lysates from naïve rats showed that all ArcNbs detect endogenous Arc as a single band (∼50 kDa) consistent with conventional anti-Arc antibodies (**Fig. 2A**). We then tested whether ALFA-ArcNbs are suitable for immunoprecipitation of endogenous, stimulus-induced Arc. CCh treatment of SH-SY5Y cells and HFS-induced LTP in the rat dentate gyrus *in vivo* are both associated with enhanced Arc expression [3, 36, 41]. SH-SY5Y cells were stimulated with 100 μM CCh for 1 h and subjected to immunoprecipitation using ALFA-ArcNb H11 and ALFA Selector^ST^ resin, followed by SDS-PAGE and immunoblotting using anti-Arc rabbit polyclonal antibody. CCh treatment induced a prominent increase in Arc expression in whole lysate, shown as input samples in figures, relative to untreated cells (**Fig. 2B**, closed arrowhead), and immunoprecipitation using ALFA-ArcNb H11 massively enriched for Arc (**Fig. 2B**, opened arrowhead). Similarly, HFS resulted in ipsilateral enhancement of Arc expression in the DG (**Fig. 2C**, closed arrowhead), which was strongly enriched in the immunoprecipitation (**Fig. 2C**, opened arrowhead). These results validate all ALFA-ArcNbs for immunoblotting and ALFA-ArcNb H11 for immunoprecipitation of endogenous Arc.

**Fig. 2.**
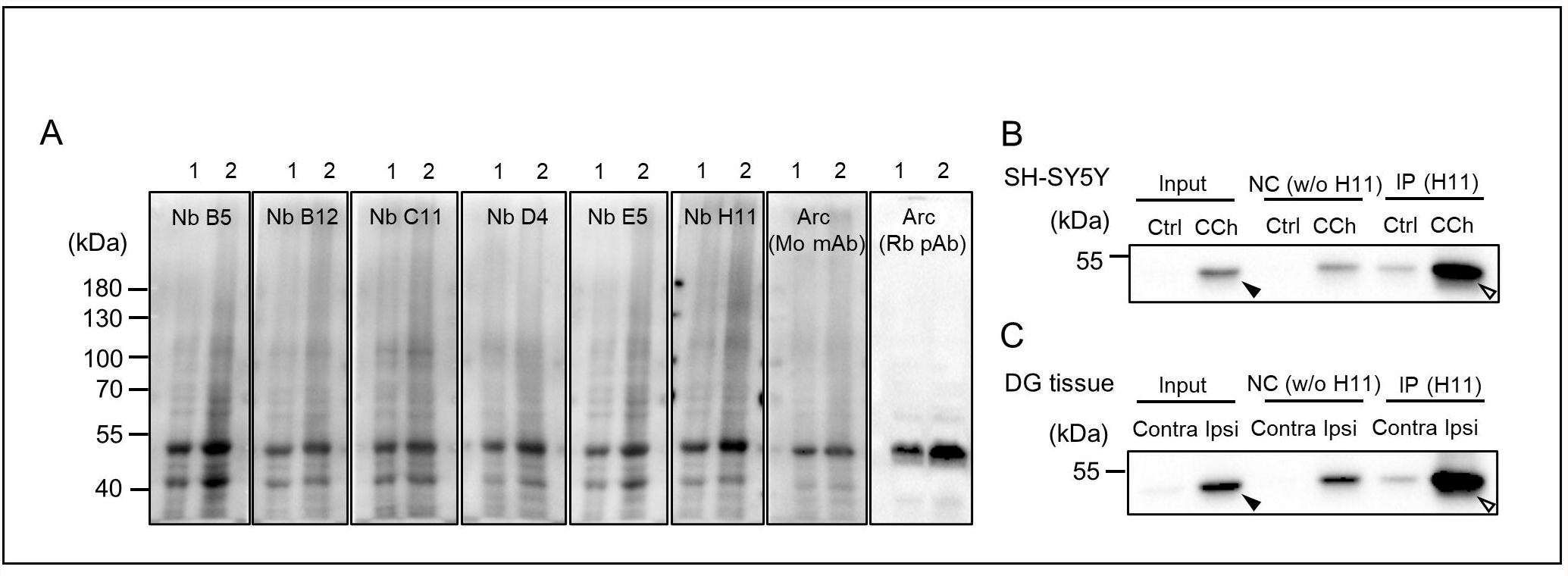
ALFA-ArcNbs for immunoblotting and immunoprecipitation analysis of endogenous Arc. ***A***, Rat cortical tissues were lysed in RIPA buffer. Different amounts of lysates were subject to SDS-PAGE followed by immunoblotting using ALFA-ArcNbs (B5, B12, C11, D4, E5, and H11) and conventional anti-Arc antibodies as reference. Immunoreactive bands corresponding to Arc were detected around 50 kDa (lane 1, 30 μg; lane 2, 60 μg). ***B***, CCh (100 µM, 1h)-treated SH-SY5Y cells were lysed and subjected to immunoprecipitation using ALFA-ArcNb H11 and ALFA Selector™ followed by SDS-PAGE and immunoblotting using anti-Arc antibody. NC, negative control (ALFA Selector^ST^ resin without ALFA-ArcNb H11. ***C***, HFS was applied unilaterally to the perforant path input to the dentate gyrus of anesthetized rats to induce LTP. At 2 hours post-HFS, DG tissue was collected from the ipsilateral (Ipsi) and contralateral (Contra) DG, and immunoprecipitation analysis was performed as described in panel B

### 3.2. ALFA-ArcNbs E5 and H11 specifically binds to Arc-NL

Next, we performed epitope mapping by expressing Arc-FL or truncation mutants in HEK293FT cells, followed by SDS-PAGE and immunoblot analysis with the six ALFA-ArcNbs. Cells were first transfected with mTq2-Arc-FL (residues 1-396, lane 1), NTR (1-140, lane 2), linker region (135-216, lane 3), and CTR (208-396, lane 4). All Nbs detected Arc-FL and CTR, but not the NTR or linker region (**Fig. 3A**). To further specific region of binding, cells were transfected with mTq2-Arc-NL (208-277, lane 5) and CL+tail (278-396, lane 6), then subjected to immunoblotting. Intriguingly, two clones E5 and H11 specifically bound to NL (**Fig. 3B**, opened arrowheads) while all other ArcNbs bound to the CL+tail region (**Fig. 3B**).

**Fig. 3.**
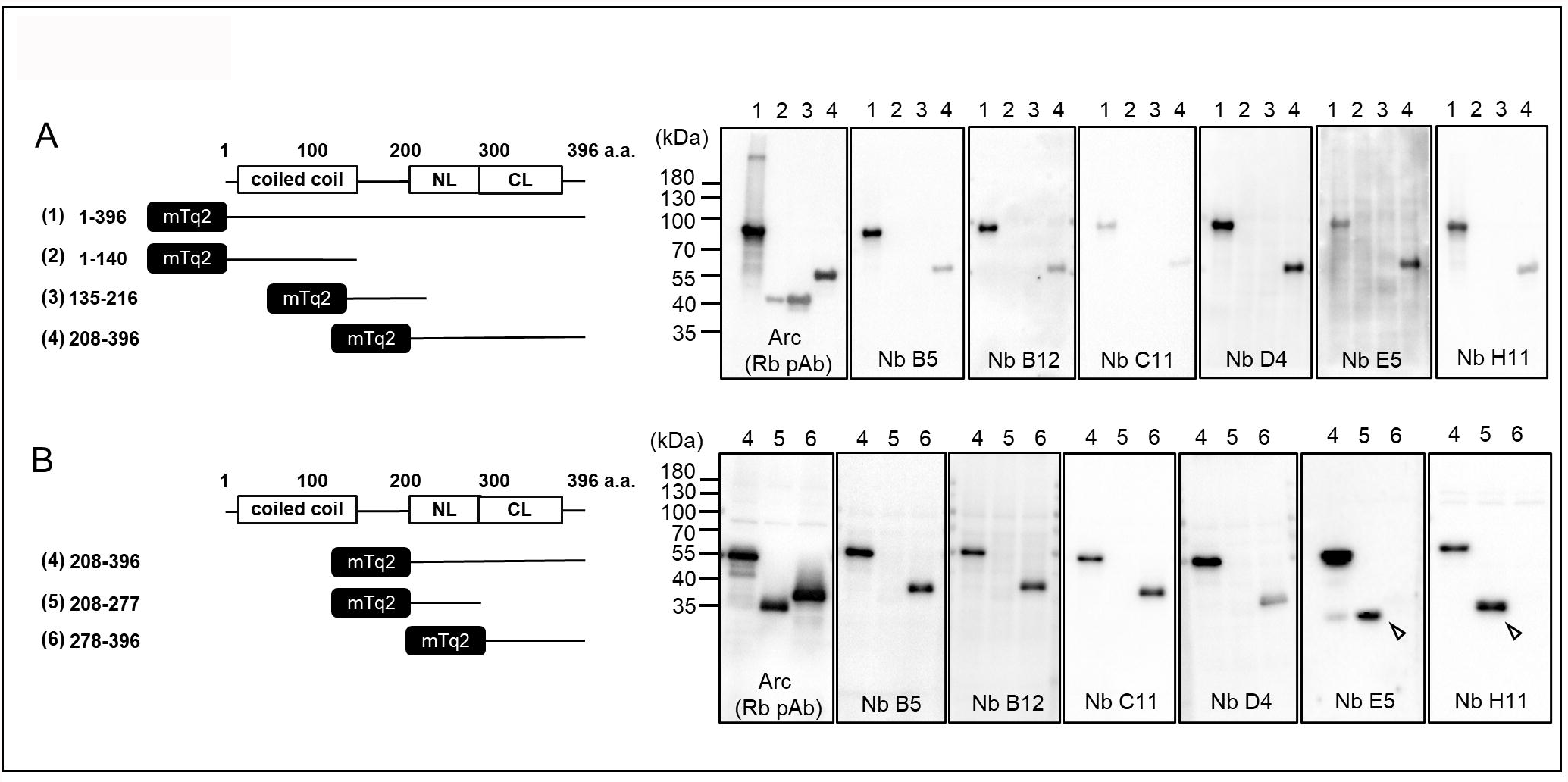
Epitope mapping of ArcNbs. ***A***, HEK293FT cells were transfected with mTq2-Arc-FL (residues 1-396) and truncation mutants expressing the Arc NTR (1-140), linker (135-216), or CTR (208-396). ***B***, Cells were transfected with mTq2-truncation mutants expressing the Arc CTR or the isolated NL and CL+tail. Cell lysates were subjected to SDS-PAGE followed by immunoblotting using ALFA-ArcNbs and anti-Arc antibody. 1, FL (residues 1-396); 2, NTR (residues 1-140); 3, linker region (residues 135-216); 4, CTR (residues 208-396); 5, NL (residues 208-277); 6, CL+tail (residues 278-370)

### 3.3. Expression of mScarlet-I-tagged and AcGFP-tagged ArcNbs as intrabodies in HEK293FT cells

We tested intracellular expression of ArcNbs in mammalian cells as genetically encoded intrabodies. HEK293FT cells were transfected with mSc-ArcNb expression vectors, then subjected to SDS-PAGE followed by immunoblotting using anti-mCh antibody to detect mSc. Expression of mSc-ArcNbs was confirmed by immunoblotting (**Fig. 4A**) and fluorescence microscopy (**Fig. 4B**). All six mSc-ArcNbs exhibited uniformly bright cytoplasmic and nucleoplasmic fluorescence without selective accumulation in subcellular compartments, formation of aggregates, or signs of morphological abnormalities. We also assessed the expression pattern of AcGFP-ArcNbs (**Fig. S3A**) AcGFP fused to either E5, C11, or H11 showed uniform fluorescence similar to the corresponding mSc-ArcNbs. However, in contrast to the mSc-ArcNbs, three of the AcGFP-ArcNbs (B5, B12, and D4) showed strong accumulation and aggregation in the nucleus (**Fig. S3B**). These results indicate that all mSc-ArcNbs and several AcGFP-ArcNbs can be expressed as intrabodies without aggregation or deleterious effects.

**Fig. 4.**
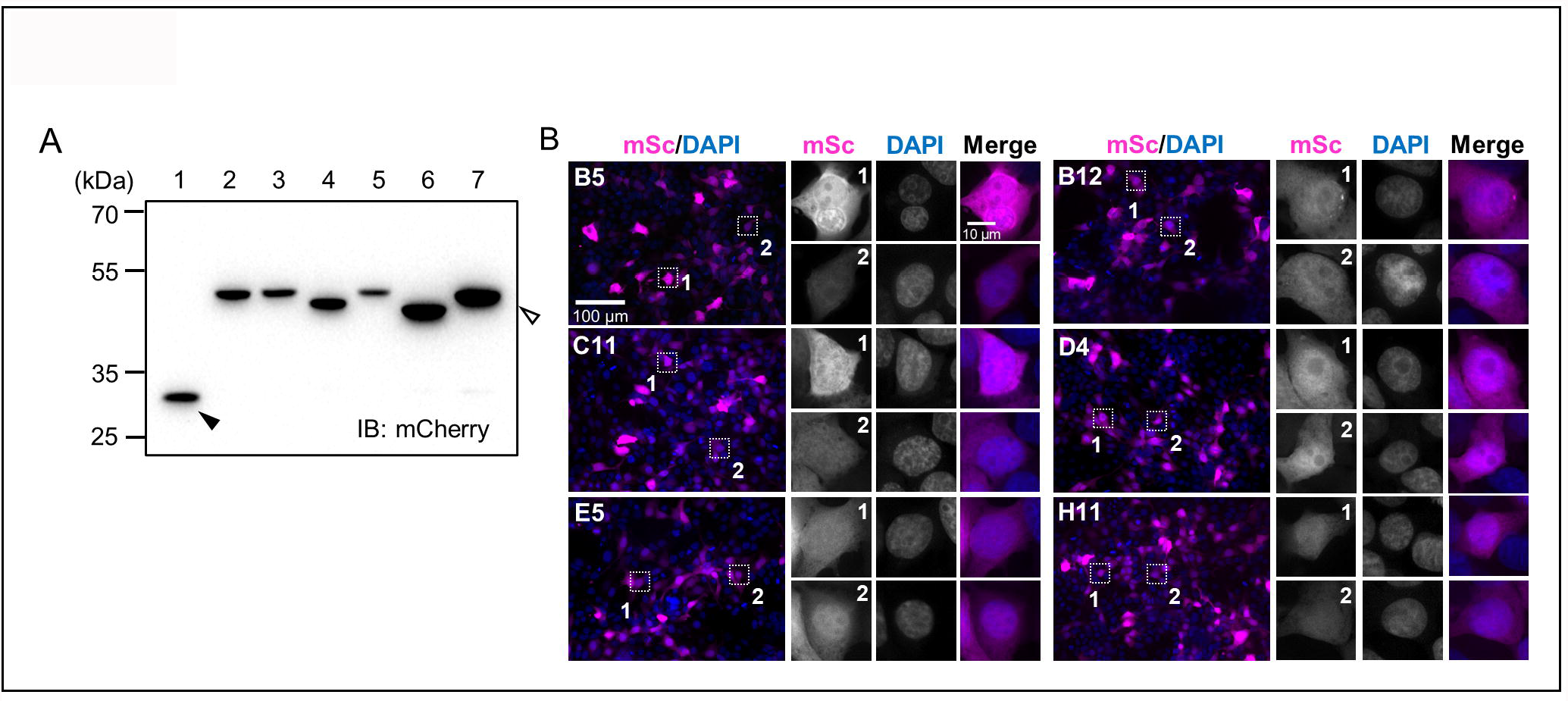
mSc-ArcNbs expression in HEK293FT cells as intrabody. ***A***, HEK293FT cells were transfected with mSc control vector (Mock, closed arrowhead) and mSc-ArcNb expression vectors (opened arrowhead). Cell lysates were subjected to SDS-PAGE followed by immunoblotting using anti-mCh antibody against mSc-tag. 1, Mock; 2, B5; 3, B12; 4, C11; 5, D4; 6, E5; 7, H11. ***B***, HEK293FT cells transfected with mSc-ArcNbs were fixed and imaged by fluorescence microscopy. Right images are high magnification images of the are in the squares in the left images. Two cells were randomly selected

### 3.4. mScarlet-I-tagged ArcNb H11 intrabody specifically binds the N-lobe and enables immunoprecipitation of intracellular Arc

We examined whether ArcNb, when expressed as intrabody, is able to bind intracellular Arc. We focused this analysis on ArcNb H11, which efficiently immunoprecipitated endogenous Arc (**Fig 2**) and selectively bound the NL in the epitope mapping analysis (**Fig. 3**). HEK293FT cells were co-expressed with mTq2-Arc (FL) together with mSc-ArcNb H11 (mSc-H11), then subjected to co-IP assay using either anti-mCh or rabbit anti-Arc polyclonal antibodies to immunoprecipitate the Arc/ArcNb complex (design illustrated in **Fig. 5A**). As expected, mSc-H11 was immunoprecipitated with anti-mCh antibody (**Fig. 5B**, closed arrowhead). Importantly, mSc-H11 in complex with mTq2-Arc (FL) was co-immunoprecipitated by anti-mCh antibody, as detected by both anti-Arc and GFP antibodies (**Fig. 5B**, opened arrowheads). In cells co-transfected with mTq2-Arc NTR or mTq2-Arc linker, no interaction of mSc-11 with these Arc regions was detected, although successful immunoprecipitation of mSc-H11 with anti-mCh antibody was confirmed (**Fig. 5C and D**, closed arrowheads). To further assess the intracellular specificity of binding, cells were co-transfected with mTq2-Arc (NL) or mTq2-Arc (CL+tail) together with mSc-H11. We found that H11 forms an intracellular complex with the Arc NL (**Fig. 5E**, opened arrowhead) but not with the CL+tail (**Fig. 5F)**. We also did a reverse co-IP assay using rabbit anti-Arc polyclonal antibody (**Fig. S4**) and found that mSc-H11 was co-immunoprecipitated with Arc FL and NL (**Figs. S4B and E**, opened arrowheads), but not with Arc NTR, linker, or CL+tail (**Figs. S4C, D and F**). Taken together, we conclude that ArcNb H11 expressed as intrabody specifically binds the Arc NL and can be used to affinity-purify intracellular full-length Arc.

**Fig. 5.**
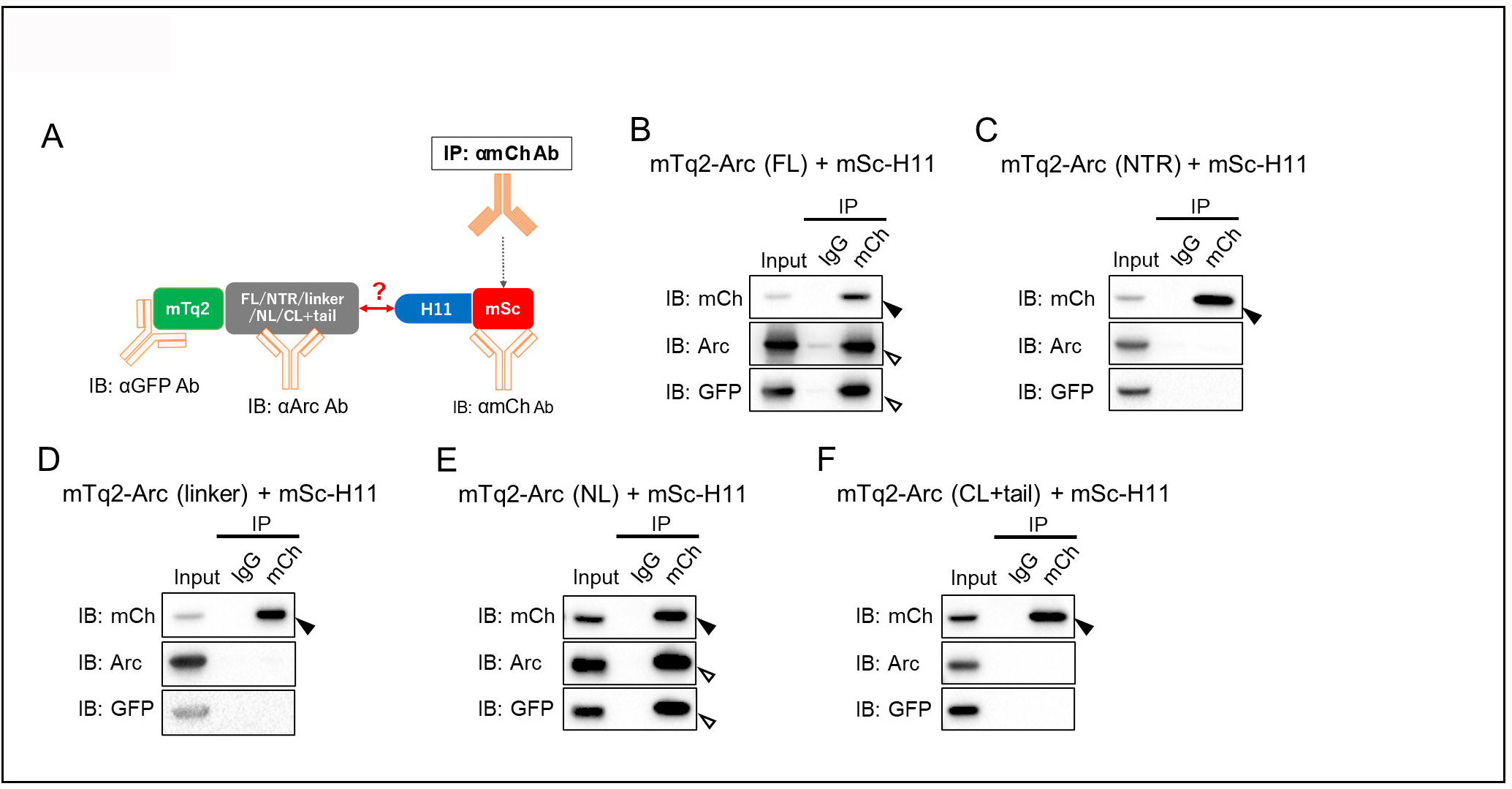
Co-immunoprecipitation of NL and FL intracellular Arc using mSc-H11 intrabody. ***A***, Schema of co-IP assay. ***B-F***, co-IP assay. HEK293FT cells were co-transfected with either mTq2-Arc-FL (***B***) or NTR (***C***) or linker region (***D***) or NL (***E***) or CL (***F***) variants and mSc-H11 constructs. Cell lysates were then subjected to co-IP assay using anti-mCh antibody. Following SDS-PAGE, immunoprecipitants were probed using anti-mCh, anti-Arc, and anti-GFP antibodies. Closed and opened arrowheads indicate immunoprecipitated mSc-H11 and co-immunoprecipitated Arc variant, respectively

## 4. Discussion

We developed and validated the first ArcNbs as new tools for probing Arc expression and function. For biochemical analyses, the small ALFA epitope tag was fused to the C-terminus of ArcNbs [33]. All clones of ALFA-ArcNbs successfully detected purified recombinant Arc protein and endogenous Arc protein from rat brain tissues (**Figs. S2 and 2A**). We show that ALFA-ArcNbs combined with anti-ALFA HRP-coupled sdAbs can be used for immunoblotting of Arc protein in the same manner as conventional anti-Arc antibodies. All six ArcNbs bound to the Arc CTR and not to the NTR or linker. Nbs E5 and H11 selectively bound to the CTD/capsid domain NL, while the remaining four Nbs bound to the segment containing the CL and C-terminal tail (**Fig. 3**).

Using a mammalian expression vector harboring mSc or AcGFP fluorescent proteins, we showed that all ArcNbs are suitable for applications as genetically encoded chromobodies (**Figs. 4A and S3B**). mSc is a bright monomeric red fluorescent protein [42, 43]. AcGFP1 is a monomeric green fluorescent protein, obtained from *Aequorea coerulescens*, with similar spectral properties to EGFP. Despite 94% amino acid sequence homology and equivalent brightness, AcGFP1 is superior to EGFP for fusion applications because it is monomeric [44] unlike EGFP that forms dimers [45, 46]. Although all mSc-ArcNbs showed uniform cytosolic and nuclear expression, three of the AcGFP-ArcNbs (B5, B12 and D4) aggregated in the nucleus (**Fig. S3B**). Further experiments are needed to understand mechanism of nuclear aggregation of these AcGFP-ArcNbs.

In the study of Markússon *et al*. (2021) on the generation and structural characterization of ArcNbs, purified untagged Nbs served as chaperones for crystallization of mammalian Arc. Atomic structures were obtained of ArcNb H11 and C11 bound to NL and CL, respectively [40]. Moreover, the complementarity determining region 3 (CDR3) of ArcNb H11 was shown to bind inside the hydrophobic pocket of the NL [40]. The CDR3 in ArcH11 harbors amino acid sequences that corresponds to the identified NL ligand binding motif [14, 20]. Isothermal titration calorimetry assays showed that ArcNb H11 binds with high affinity (1 nM K_d_) and competitively displaces stargazin, an auxiliary subunit of AMPA-type glutamate receptor, which has the highest binding affinity (34.9 μM) of the known Arc-NL ligands [40]. The present work shows that genetically encoded ArcNb H11 expressed in mammalian cells binds specifically to the NL and enables co-immunoprecipitation of intracellular Arc (**Figs. 5 and S3**). ArcNb H11 also enabled efficient immunoprecipitation of endogenous Arc synthesized during the maintenance phase of LTP in the rat dentate gyrus *in vivo* (**Figs. 2**). Taken together, these findings make ArcNb H11 attractive as a high-affinity binder for labeling and tracking Arc inside cells, and a candidate competitive inhibitor of Arc-NL function *in vivo*.

ArcNbs B5, B12, C11, and D4 were shown to bind to the Arc region encompassing the CL and C-terminal tail (**Fig. 3B)**. However, specific association with the CL is likely, as recombinant purified ArcNbs bound to the isolated CTD (Markússon *et al*. (2021). Furthermore, crystal structure analysis of untagged ArcNb C11 in complex with the CTD demonstrated binding to the CL in a region that encompasses a conserved retroviral capsid dimerization motif [14, 24, 40]. Thus, ArcNb C11 is of interest as a potential regulator of CL-mediated dimerization in the context of full-length Arc oligomerization, and eventual capsid assembly. Although CL is structurally homologous to NL [14], there is no evidence of similar peptide ligand binding [20, 25]. The Arc C-terminal tail (370-396) has a PEST signal region (a sequence segment that is rich in P, E, S, T residues) [47] implicated in Arc degradation [48].

In conclusion, we developed two lines of ArcNbs: one (clones E5 and H11) for binding to NL, another (clones B5, B12, C11, and D4) for binding the CL and C-terminal region. These new tools open a range of possibilities, including expression of intrabodies for labelling and tracking endogenous Arc in live-cell imaging experiments and for biochemical isolation of native Arc complexes. Given their stable expression and high-affinity binding, it will be important in future work to evaluate ArcNbs as genetically-encoded inhibitors of Arc NL signaling and oligomerization in neurons. ArcNbs can be used and further developed to elucidate Arc domain-specific mechanisms and functions in synaptic plasticity. The recombinant ALFA-tagged ArcNbs may also be useful in efforts to detect oligomers and isolate intact capsids from brain tissue and other sources.

## CRediT author statement

**Yuta Ishizuka:** Conceptualization, Methodology, Validation, Investigation, Resources, Data Curation, Writing – Original Draft, Visualization, Supervision, Project administration. **Tadiwos F. Mergiya:** Validation, Investigation, Data Curation, Writing – Review & Editing. **Rodolfo Baldinotti:** Investigation, Validation. **Ju Xu:** Investigation, Resources. **Erik I. Hallin:** Resources. **Sigurbjörn Markússon;** Resources, **Petri Kursula:** Conceptualization, Resources, Project administration. Supervision, Funding acquisition. **Clive R. Bramham:** Conceptualization, Writing – Review & Editing, Supervision, Methodology, Resources, Data Curation, Funding acquisition, Project administration.

## Declarations

### Ethical Approval

All experimental procedures were approved by the Norwegian National Research Ethics Committee in compliance with EU Directive 2010/63/EU, ARRIVE guidelines. All methods were carried out in accordance with relevant guidelines and regulations. Persons involved in the animal experiments have approved Federation of Laboratory and Animal Science Associations (FELASA) C course certificates and training.

### Consent for Publication

All authors reviewed and approved the manuscript.

## Data availability

The datasets generated and analyzed during the current study are available from the corresponding authors on reasonable request.

## Acknowledgements

We would like to thank The Laboratory Animal Facility of University of Bergen for maintaining experimental animals. This work was supported by a Research Council of Norway Toppforsk grant (249951) to CRB.

## Supplemental Materials

**Fig. S1**. Quality analysis of purified ALFA-ArcNbs. All purified ALFA-ArcNbs were subjected to SEC followed by SDS-PAGE (***A***, B5; ***B***, B12; ***C***, C11; ***D***, D4; ***E***, E5; ***F***, H11). Upper images are acryl amid gel stained by InstantBlue® Coomassie Protein Stain (Abcam plc, Cambridge, UK) after SDS-PAGE. Opened arrowheads indicate purified ALFA-ArcNbs around 15 kDa. Lower images are SEC chromatograms. Closed arrowheads indicate the fractions which were collected.

**Fig. S2**. All ArcNbs detect purified Arc recombinant protein both in native and denatured condition. ***A***, Purified rat Arc mutant protein (Arc^s113-119A^) was subjected to native-PAGE followed by immunoblotting using ALFA-ArcNbs and conventional anti-Arc antibodies. BSA was used as reference for PAGE (∼66 kDa in Ponceau S staining). Std., Molecular weight marker; 1, BSA (1 μg); 2, rArc mutant protein (0.1 μg); 3, rArc mutant protein (0.5 μg). ***B***, Purified Arc^s113-119A^ was subjected to SDS-PAGE followed by immunoblotting. 4, rArc mutant protein (0.05 μg); 5, rArc mutant protein (0.1 μg); 6, rArc mutant protein (0.5 μg).

**Fig. S3**. AcGFP-ArcNbs expression in HEK293FT cells as intrabody. ***A***, HEK293FT cells were transfected with AcGFP control vector (Mock, closed arrowhead) or AcGFP-ArcNb expression vectors (opened arrowhead). Cell lysates were subjected to SDS-PAGE followed by immunoblotting using anti-GFP antibody. 1, Mock; 2, B5; 3, B12; 4, C11; 5, D4; 6, E5; 7, H11. ***B***, HEK293FT cells transfected with AcGFP-ArcNbs were fixed and imaged by fluorescence microscopy. Right images are high magnification images of the area in the squares in the left images. Two cells were randomly selected.

**Fig. S4**. Reverse co-IP assay using anti-Arc antibody. ***A***, Schema of Co-IP assay. ***B-F***, Co-IP assay. HEK293FT cells were co-transfected with either mTq2-Arc-FL (***B***) or NTD (***C***) or Linker (***D***) or NL (***E***) or CL (***F***) variants and mSc-H11 constructs. Cell lysates were then subjected to co-IP assay using anti-Arc antibody. Following SDS-PAGE, immunoprecipitants were probed using anti-mCh, Arc, and GFP antibodies. Closed and opened arrowheads indicate immunoprecipitated mTq2-Arc variants and co-immunoprecipitated mSc-H11, respectively.

