## Supplemental Figures for "Development and validation of Arc nanobodies: new tools for probing Arc dynamics and function"

### Fig. S1 (A-D)

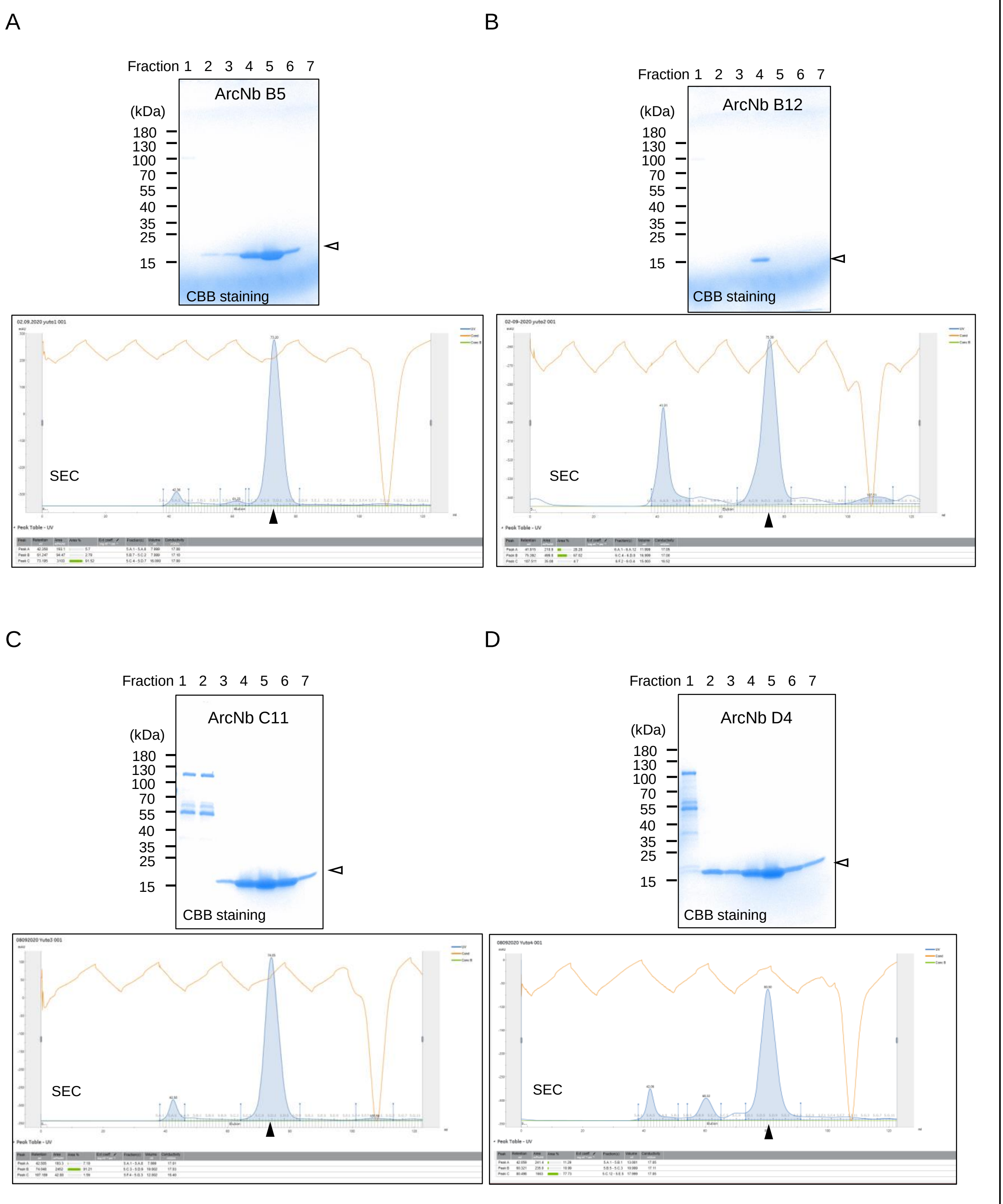


174 mm (double column) x 234 mm (Max)

#### Fig. S1 (E&F)

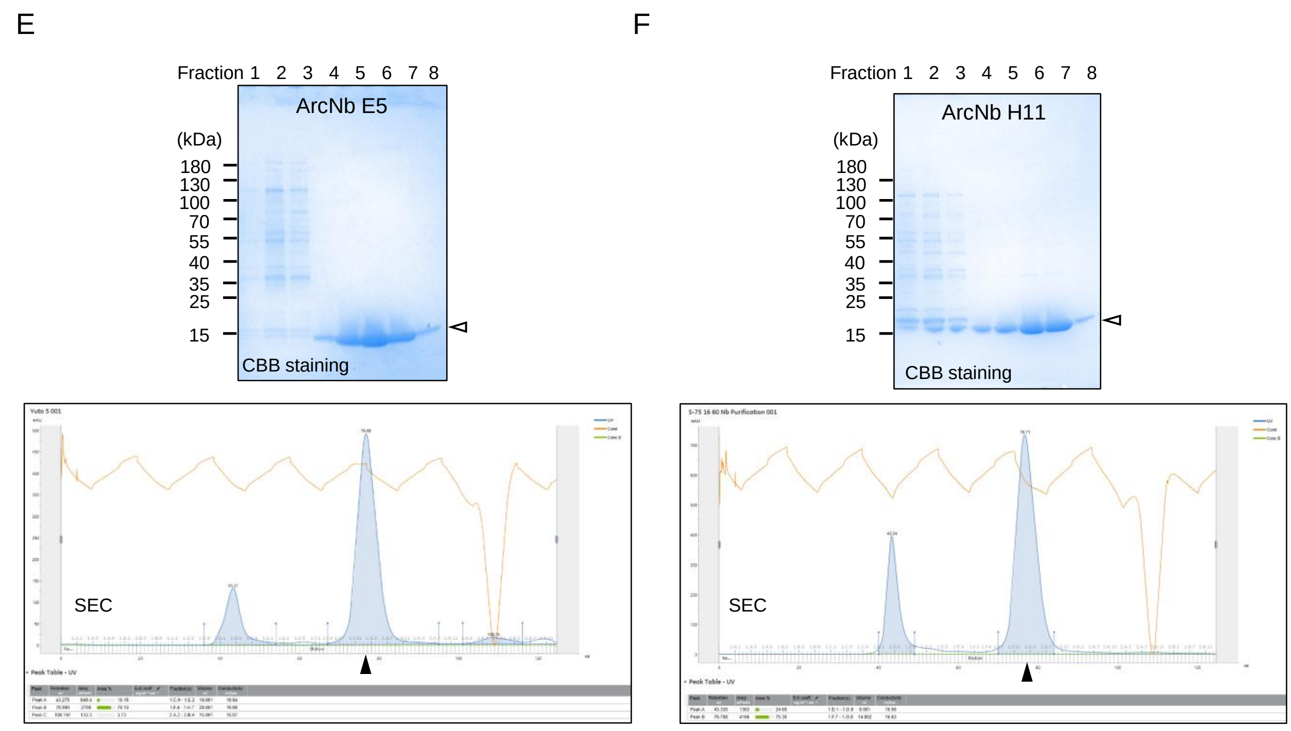


174 mm (double column)

#### Fig. S2

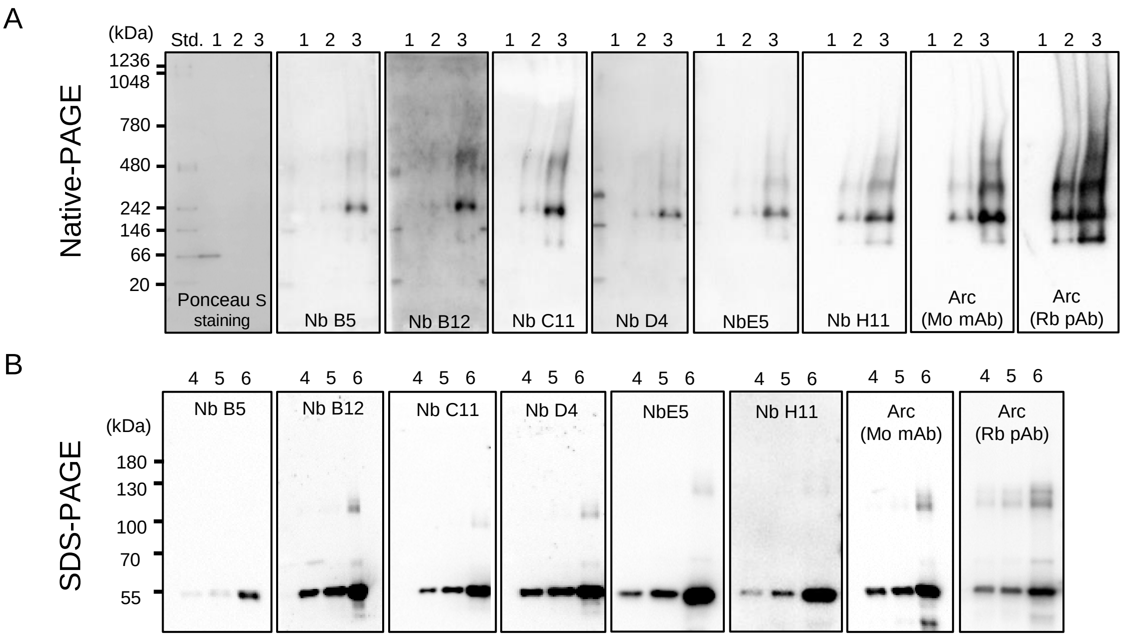


174 mm (double column)

#### Fig. S3

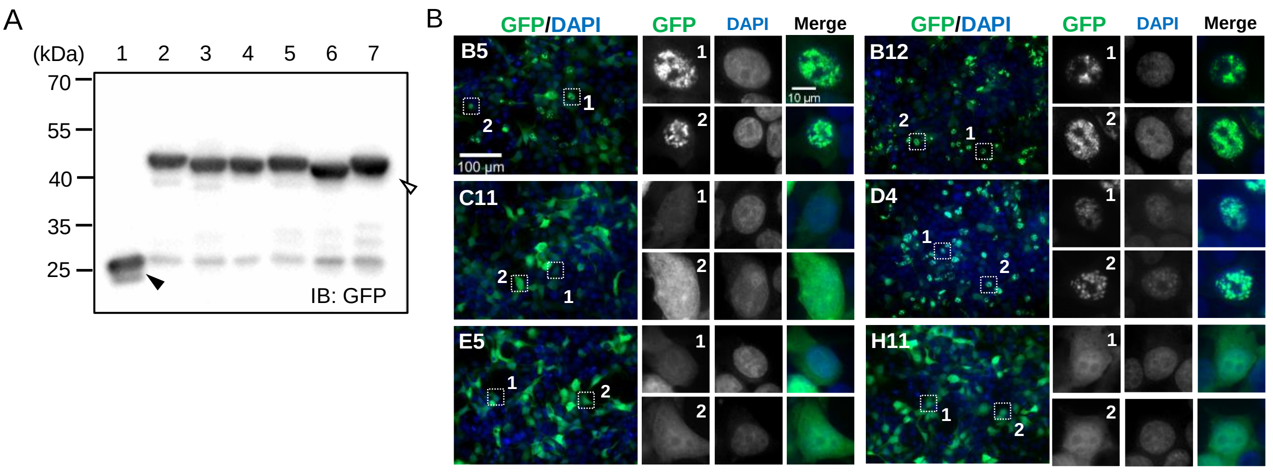


174 mm (double column)

#### Fig. S4

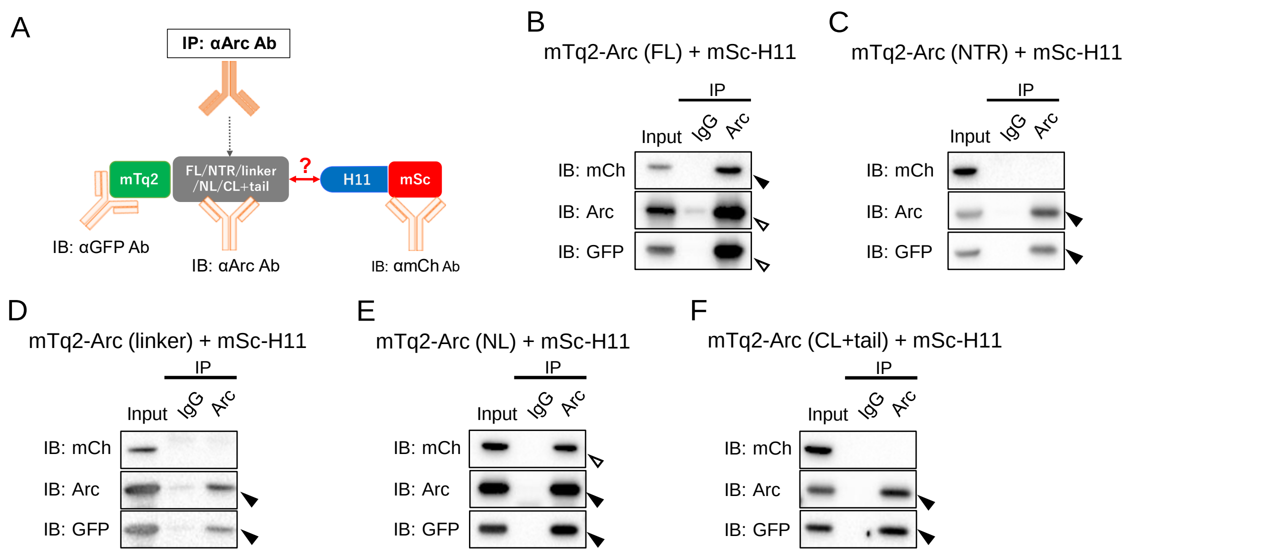


174 mm (double column)
